# Comparison of Transcriptomic Changes in Younger and Older Multiple Myeloma Patients from the MMRF-CoMMpass Study RNA-Seq Data

**DOI:** 10.1101/2022.09.14.507921

**Authors:** Merve Keskin

## Abstract

**Background:** Age-related differences in Multiple Myeloma (MM) are studied in clinical and genomic context, however, transcriptome changes have not yet been determined. The aim of this study is to identify the genes that are expressed differently in young and old patient groups and to examine the relationship of these genes with biological pathways and the drugs that can be used.

**Methods:** The MMRF CoMMpass cohort RNA-Seq data (n=634) was used to analyze differentially expressed genes between young and old patients. GO term and KEGG gene-set enrichment analysis were conducted using R packages. Drug-gene interactions were detected using DGIdb.

**Results:** Globally, 523 genes (366 upregulated, 157 downregulated) were differentially expressed (p < 0.05) in young patients. Totally 220 GO terms, mostly related to immune regulation pathways were enriched. “Cytokine-cytokine receptor interaction” gene-set was enriched in KEGG GSEA. Among the highest expression difference, genes involved in immune regulation (FCGR1A, FCER1G, TLR2), known proto-oncogenic genes (BCL2, FGR) and genes under investigation for association with various cancers (RGL4, MT-RNR1, ETS2, ENPP3, FUT7, NTNG2, PRAM1) were identified. Drugs associated with the pathways affected by these genes were identified.

**Conclusions:** Further investigation of differentially expressed genes in young patients may shed light on new treatment options.

## 1. Introduction

Multiple myeloma (MM) is a malignancy of bone marrow terminally differentiated plasma cells, a disease mostly affects the elderly, diagnosed at a median age approximately 66–70 years and characterized by diverse and complex tumor genetics. It is extremely rare in those less than 30 years of age with a reported frequency of 0.02% to 0.3% (Kazandijan 2016).

Aging is a risk factor for many diseases and cancer is one of these, however the age-related biological characteristics of cancer are still not elucidated at the molecular level. Comparing the clinical and genetic features of patients presenting at an early age with the presentation of older age is important to better understand the pathogenesis of the disease and to understand whether it is necessary to evaluate different treatment options in different age groups (Shah et al. 2021).

In a recent study that analyzed multiple cancer genome, transcriptome and methylome data from the TCGA (The Cancer Genome Atlas) cohort to understand the biology of cancers in younger versus older individuals, six TCGA tumors showed an age-associated outcome and molecular landscape. The study also demonstrated that aging-associated dysfunction is tumor type specific and suggested that could lead to different therapeutic strategies (Shah et al. 2021).

In another comprehensive study, using the TCGA dataset with a multi-omics approach, the largest age-related genomic differences were found in gliomas and endometrial cancer, and the study identified age-related global transcriptomic changes that were partly regulated by age-associated DNA methylation changes (Chatsirisupachai et al. 2021).

Independent studies comparing clinical features of younger and older MM patients have reported varying results, with some reporting no difference in presenting features of young and old MM patients (Corso et al. 1998), while other studies reported bone lytic lesions, renal impairment, hypercalcemia, light-chain myeloma and extramedullary disease are higher in younger patients (Duek et al. 2021),(Jurczyszyn et al. 2016) (Pydi et al. 2021). Most studies concluded that survival was significantly better in the younger patients compared with older patients patients (Jurczyszyn et al. 2016) (Boyle et al. 2020) (Ludwig et al. 2010) (Pál, Illés, and Váróczy 2020), although one of the studies claimed that this difference was lost when adjusted for differences in life expectancy (Corso et al. 1998).

Age related cytogenetic anomalies in MM were studied in a large cohort study comparing young and elderly patient groups and conventional cytogenetic analysis showed no difference in the frequency of any cytogenetic abnormality (Ludwig et al. 2008), however, another study reported that translocation t(11; 14) was more prevalent in younger myeloma patients and it is related to poor outcome (Duek et al. 2021). Another study investigating mutational signatures showed that mutational signatures did not substantially differ between age groups, except for the higher rate of APOBEC (SBS2 and 5) in the group >80 years and older and also found that simple and complex structural variants and the prevalence of chromothripsis increased with age (Boyle et al. 2020).

However, there is no information available yet about the differences between young and old MM patients at transcriptome level, which is important to understand the possible different pathogenic processes and determine the need for different treatment strategies.

The aim of this study is to compare the RNA-Seq data of the MMRF (Multiple Myeloma Research Foundation) CoMMpass study in young and old patient groups, to identify differentially expressed genes and examine the biological pathways and drug interactions associated with these genes.

## 2. Materials and Methods

### 2.1 Data Collection

All data in this study were generated as part of the Multiple Myeloma Research Foundation (MMRF) Personalized Medicine Initiatives (https://research.themmrf.org and www.themmrf.org). The MMRF CoMMpass cohort (NCT01454297) includes 1143 multiple myeloma patients from 84 clinical centers located in the United States, Canada, Spain, and Italy; in which tumor samples were characterized using whole genome, exome, and RNA sequencing at diagnosis and progression, and clinical data was collected every three months. The cohort was conducted in accordance with the Declaration of Helsinki and all patients provided written informed consent.

In this study, 634 newly diagnosed MM patients from CoMMpass study, whose bone marrow sample CD138+ cells RNA-Seq data available were included. RNA-Seq STAR method unstranded counts of 634 patients in the IA17 cohort of the MMRF CoMMpass trial were employed to detect DEGs.

The RNA-seq and clinical data were downloaded on 1 July 2022, the source files “MMRF_CoMMpass_IA17_star_geneUnstranded_counts.tsv” and “MMRF_CoMMpass_IA17_PER_PATIENT_V2.tsv” are available upon request from https://research.themmrf.org.

### 2.2 Study Design

The study design and the workflow of this study is summarized in Figure 1.

**Figure 1.**
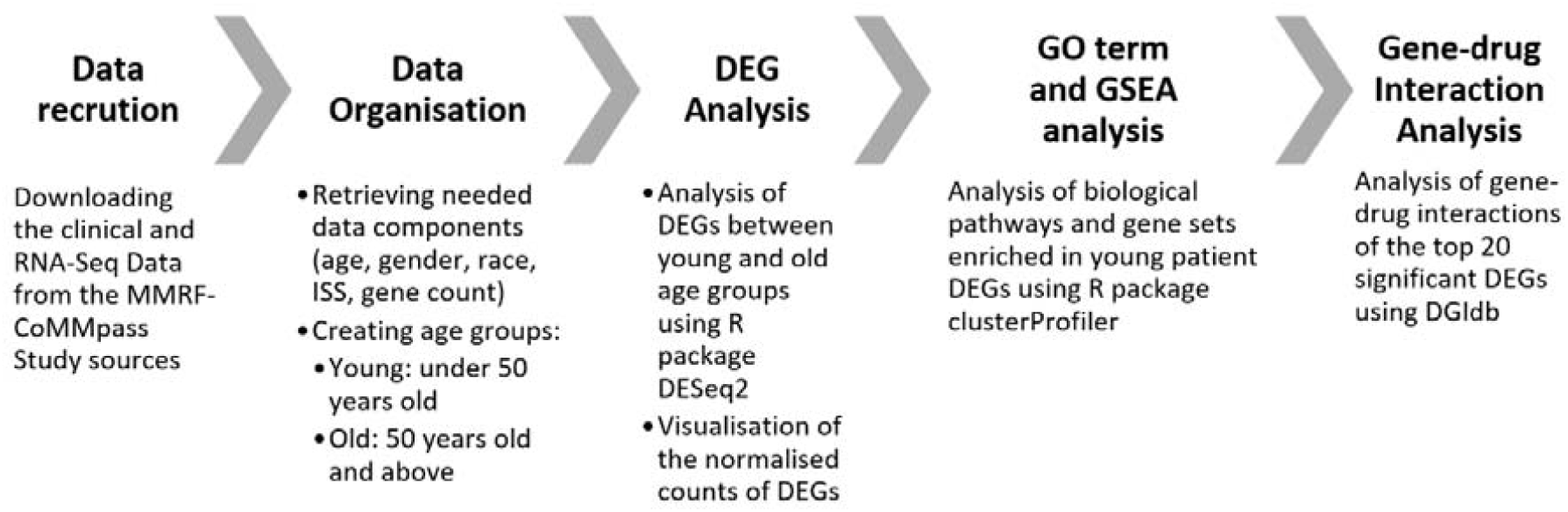
The study design and workflow of the study. (DEG: differentially expressed gene, DGIdb: Drug-Gene Interaction Database ISS: International Staging System, GO: Gene ontology, GSEA: Gene-set Enrichment Analysis, MMRF: Multiple Myeloma Research Foundation)

All R codes used to analyze the data in this study are available at https://github.com/merrvve/commpas-deg.

### 2.3 Count Normalization

All statistical analyses in this study were performed with the statistics software R (version 4.2.1; available from: https://www.r-project.org) and R packages developed by BioConductor project (available from: https://www.bioconductor.org). The expression level of each gene was summarized and normalized using the DESeq2 R/Bioconductor package which by default uses the median of ratios method (Love MI, Huber W 2014).

### 2.4 Differential Gene Expression Analysis

Differential gene expression analyses were performed using DESeq2 pipeline (Love MI, Huber W 2014). p values were adjusted with the default option of the DESeq2 package. Genes were considered differentially expressed if they had an adjusted p value □<□0.05 and a |log2 fold change|> 0.58.

### 2.5 Gene Ontology and Gene Set Enrichment Analysis

Gene Ontology (GO) pathway and Kyoto Encyclopedia of Genes and Genes (KEGG) gene-set enrichment analysis (GSEA) were performed using R package clusterProfiler (G Yu, LG Wang, Y Han 2012) to investigate the biological function and signaling pathway of DEGs, with a p value <0.05 as cut-off value for significance.

### 2.6 Gene-Drug Interaction Analysis

The interactions of the top 20 significant DEGs with prescription drugs were identified using the Drug Gene Interaction Database (DGIdb) v4.2.0 (https://www.dgidb.org access 13.08.2022) (Freshour et al. 2021). date:

## 3. Results

### 3.1 Patient Characteristics

A total of 634 patients in the MMRF CoMMpass cohort with bone marrow CD138+ cells RNA-Seq data were included in this study. Patients were aged 45–93 years (median 64) and were categorized as under 50: young or 50 years and over: old. The characteristics of the patient groups are summarized in Table 1.

**Table 1.**
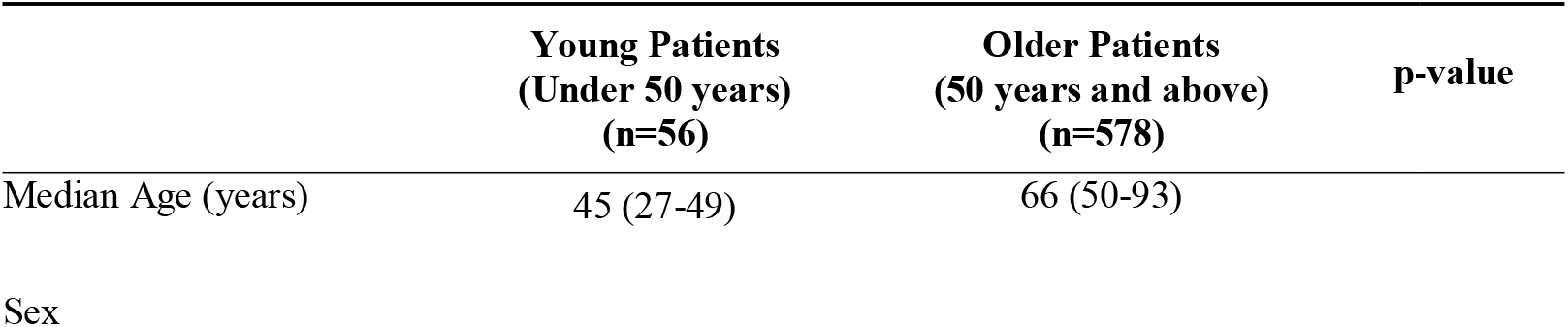
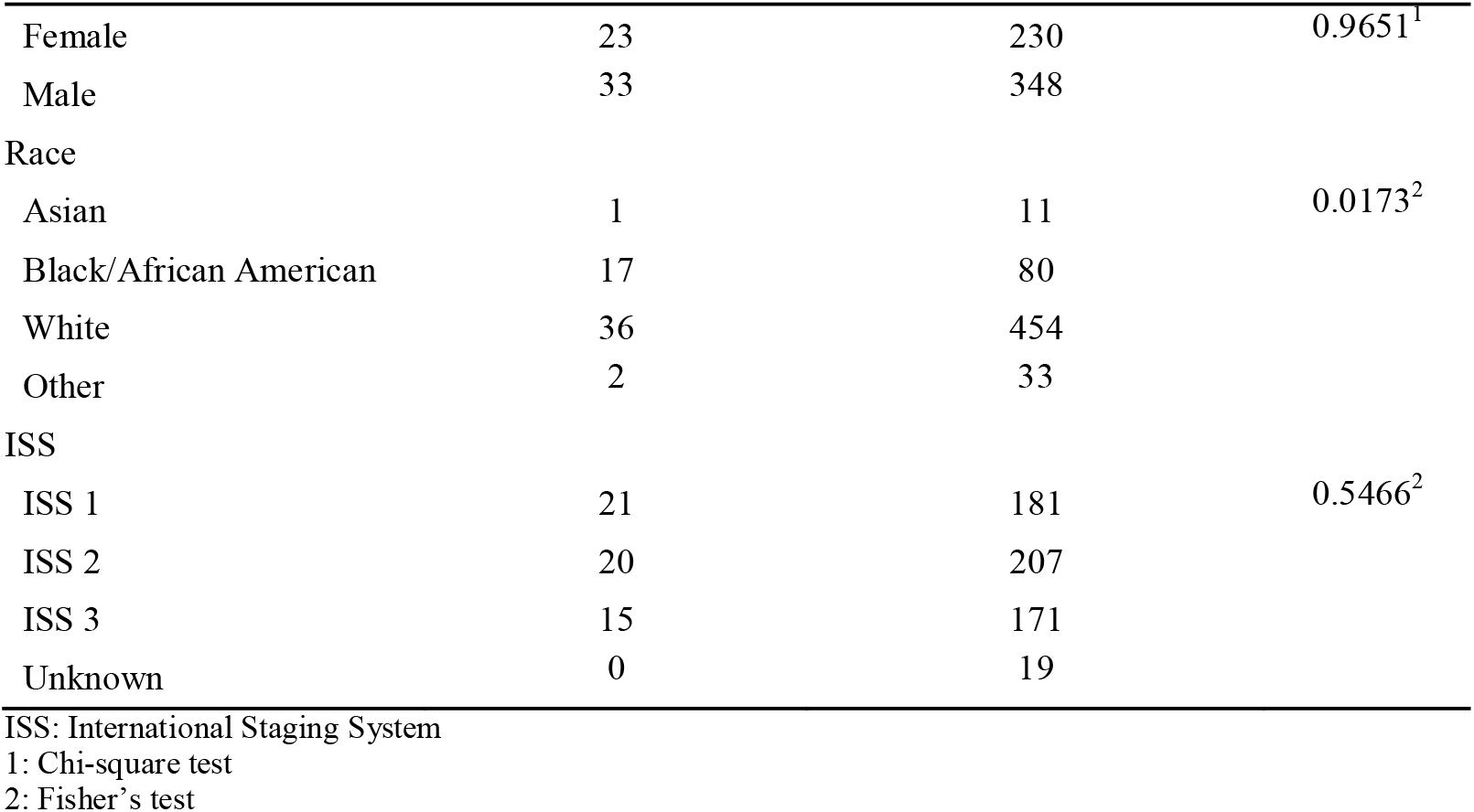
Patient Characteristics (n=634)

|  | Young Patients<br>(Under 50 years)<br>(n=56) | Older Patients<br>(50 years and above)<br>(n=578) | p-value |
| --- | --- | --- | --- |
| Median Age (years) | 45 (27-49) | 66 (50-93) |  |
| Sex |  |  |  |
| Female | 23 | 230 | 0.9651<sup>1</sup> |
| Male | 33 | 348 |  |
| Race |  |  |  |
| Asian | 1 | 11 | 0.0173<sup>2</sup> |
| Black/African American | 17 | 80 |  |
| White | 36 | 454 |  |
| Other | 2 | 33 |  |
| ISS |  |  |  |
| ISS 1 | 21 | 181 | 0.5466<sup>2</sup> |
| ISS 2 | 20 | 207 |  |
| ISS 3 | 15 | 171 |  |
| Unknown | 0 | 19 |  |

ISS: International Staging System
1: Chi-square test
2: Fisher's test

### 3.2 Differential Gene Expression

Globally, RNA-Seq differential gene expression analysis showed 523 genes differentially expressed with p < 0.05 and |Log2 FC| > 0.58 in young patients, relative to older patients. Overall, 366 of them were upregulated and 157 were downregulated. Normalized RNA-Seq counts of all genes with labeled DEGs are shown in Figure 2. Full list of the significant DEGs is included in Supplementary Data 1.

**Figure 2.**
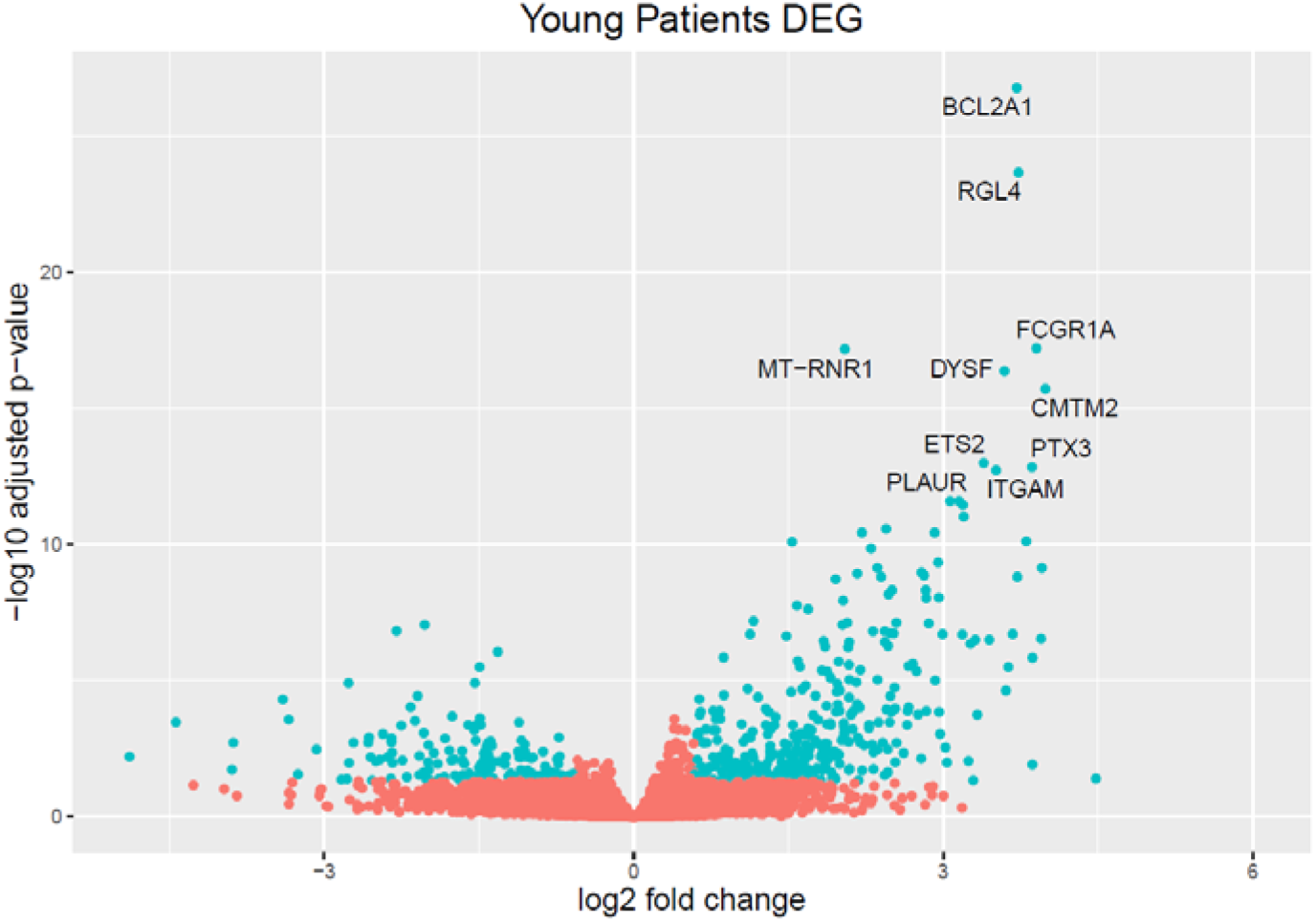
Volcano Plot of Differently Expressed Genes (DEG) in Young Patient Group Relative to Old Patients. DEGs in younger patients are colored in green and top 10 DEGs are labeled with gene symbols.

Top 20 significant DEGs ranked by adjusted p values were all upregulated in younger patients. The plot of normalized counts of the top 20 DEGs is shown in Figure 3. Among these genes, *BCL2A1 and FGR* were proto-oncogenes; *FCGR1A, FCER1G and TLR2* were related to immune regulation. The description and possible relation to oncogenic pathways of the top 20 significant DEGs are summarized in Table 2.

**Table 2.**
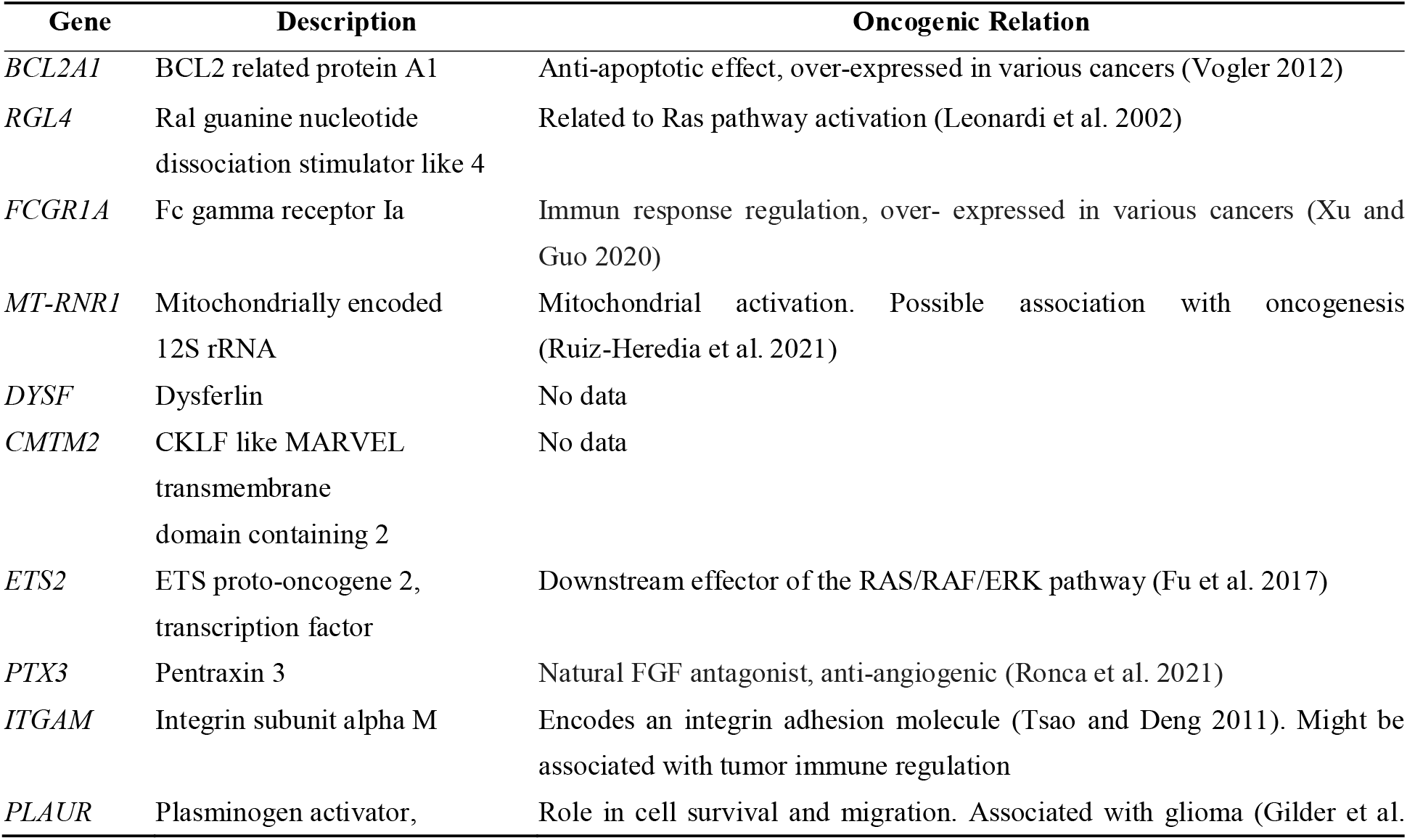
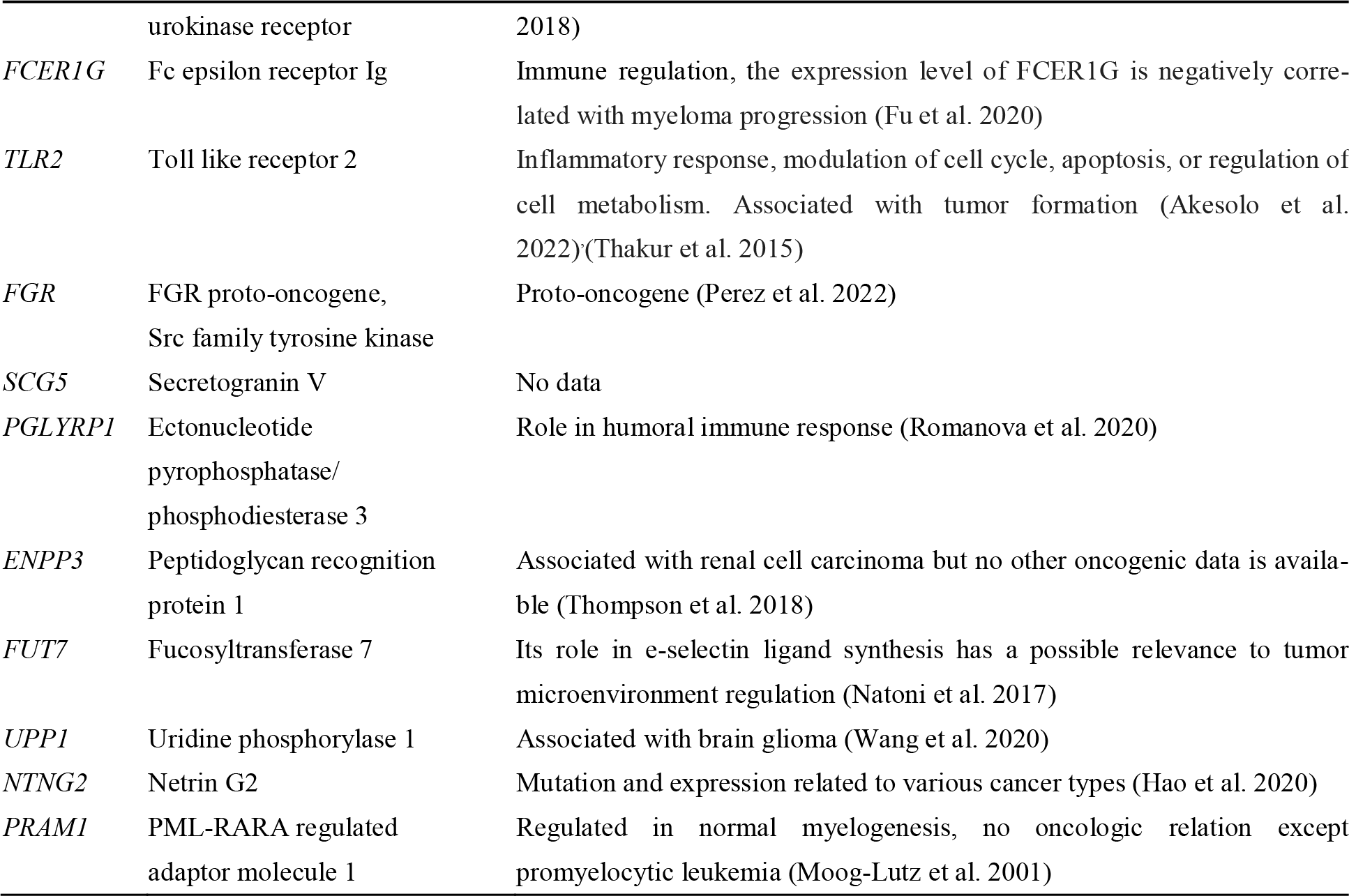
Description of the Top 20 DEGs in Young Patients.

**Figure 3.**
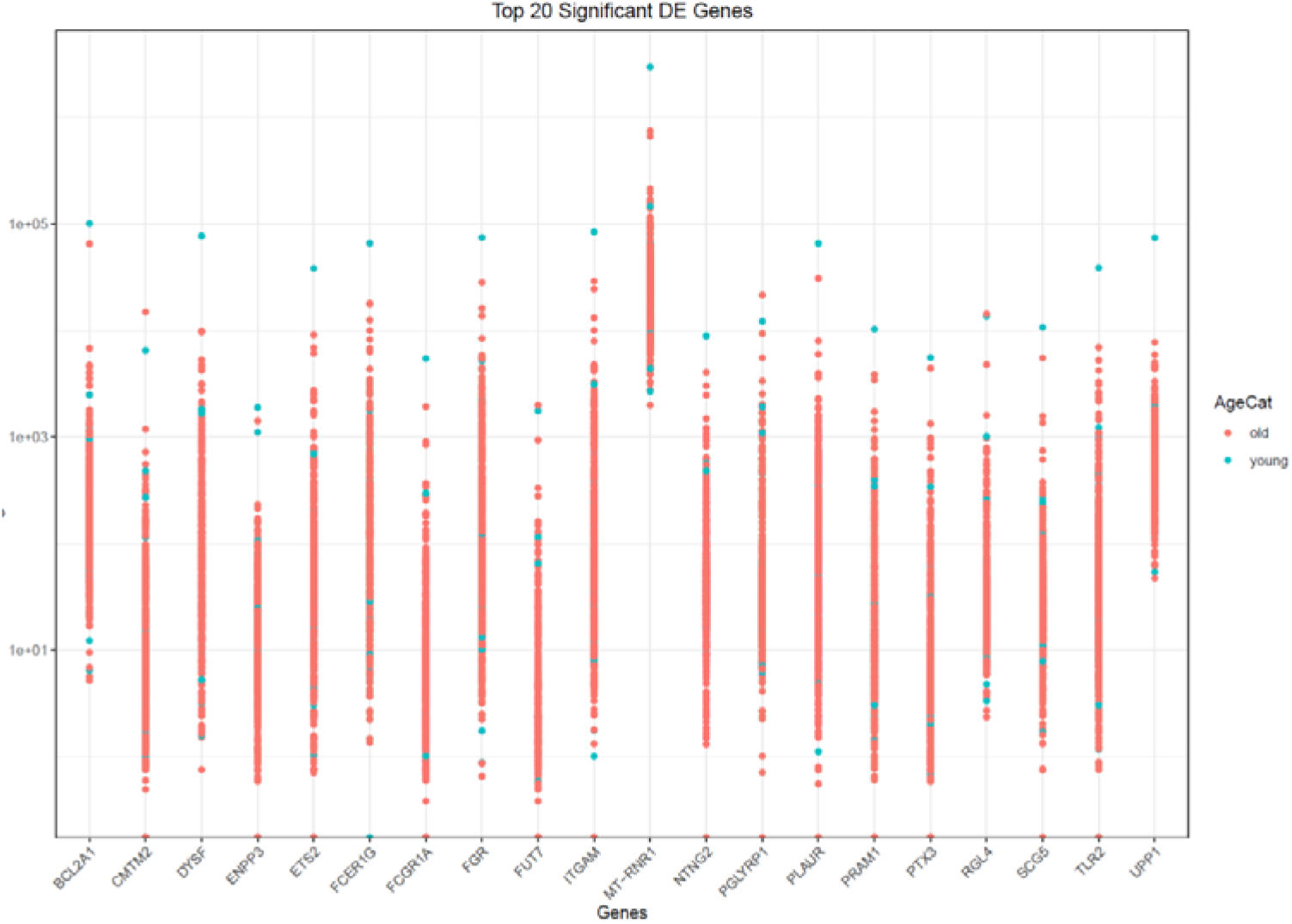
Normalized Counts of Top 20 Significant DEGs in Young Patients Group

### 3.3 GO Term and KEGG Analysis

Further analysis of KEGG gene-set enrichment and GO term enrichment were performed to find the overview functions of differentially expressed genes. GO enrichment analysis revealed that significant pathways were mostly associated with immune regulation. Totally, 220 GO biological pathway terms were enriched with adjusted p values between 3.78 × 10^−8^ – 0.04, and q values between 3.29 × 10^−8^ – 0.04. The top 50 significant GO terms are shown in Figure 4. Full list of significant GO terms with p and adjusted p values are included in Supplementary Data 2. Among these, the top five significant GO terms were leukocyte migration, myeloid leukocyte migration, T cell activation, regulation of lymphocyte activation and cytokine-mediated signaling pathway. The category net plot showing the related DEGs with these pathways are summarized in Figure 5.

**Figure 4.**
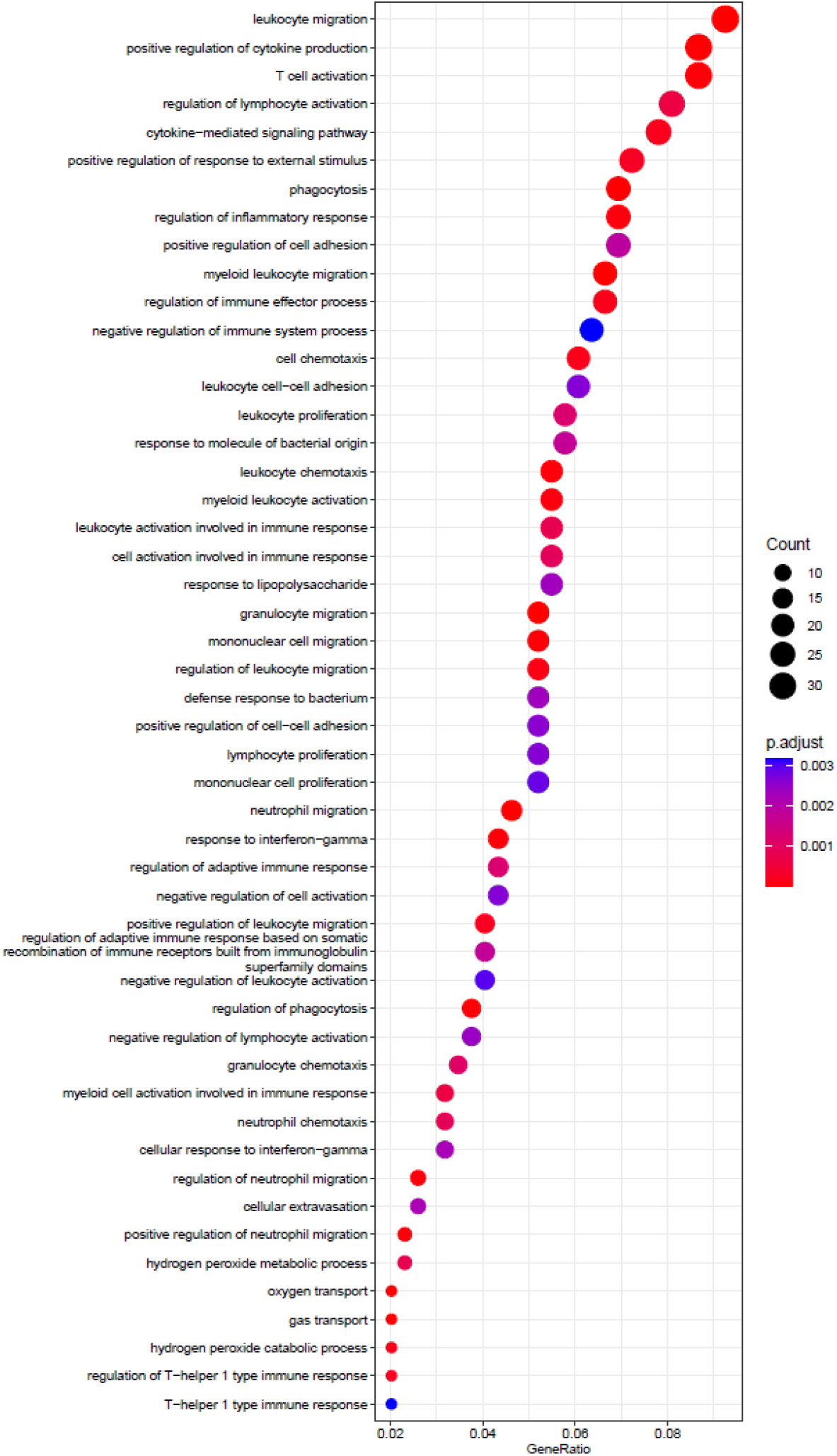
The number of genes associated with the first GO 50 terms: GO terms (size) and the p-adjusted values for these terms (color)

**Figure 5.**
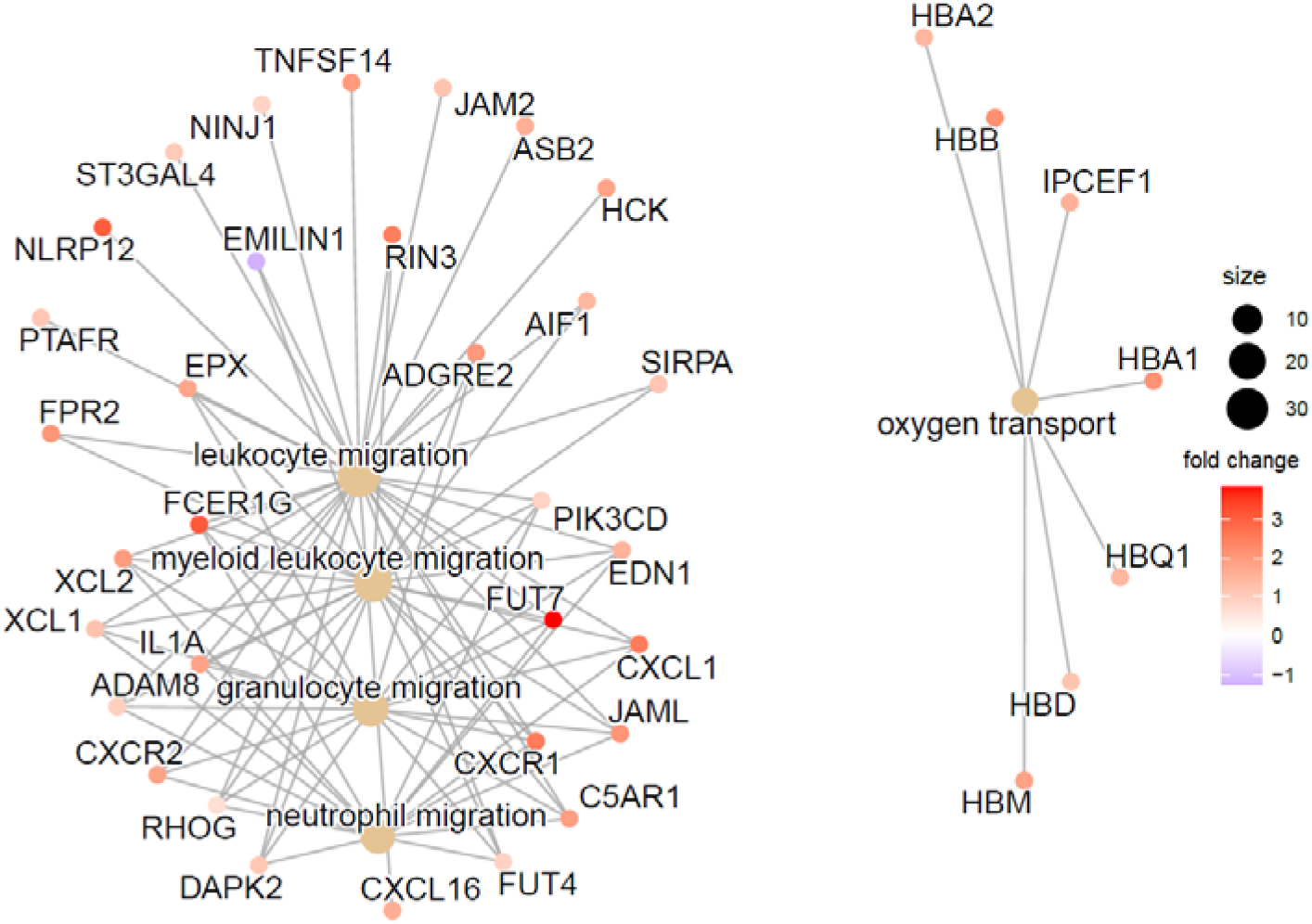
Category Net Plot of the top five significant GO terms and the fold changes of the significant genes associated with these terms.

By KEGG gene set enrichment analysis (GSEA), only one gene set, “cytokine-cytokine receptor interaction” (hsa04060, available from https://www.genome.jp/pathway/hsa04060) pathway was enriched (adjusted p value 0.01 and q value 0.007) in young MM patient DEGs. GSEA plot of the enriched cytokine-cytokine receptor interaction pathway is shown in Figure 6.

**Figure 6.**
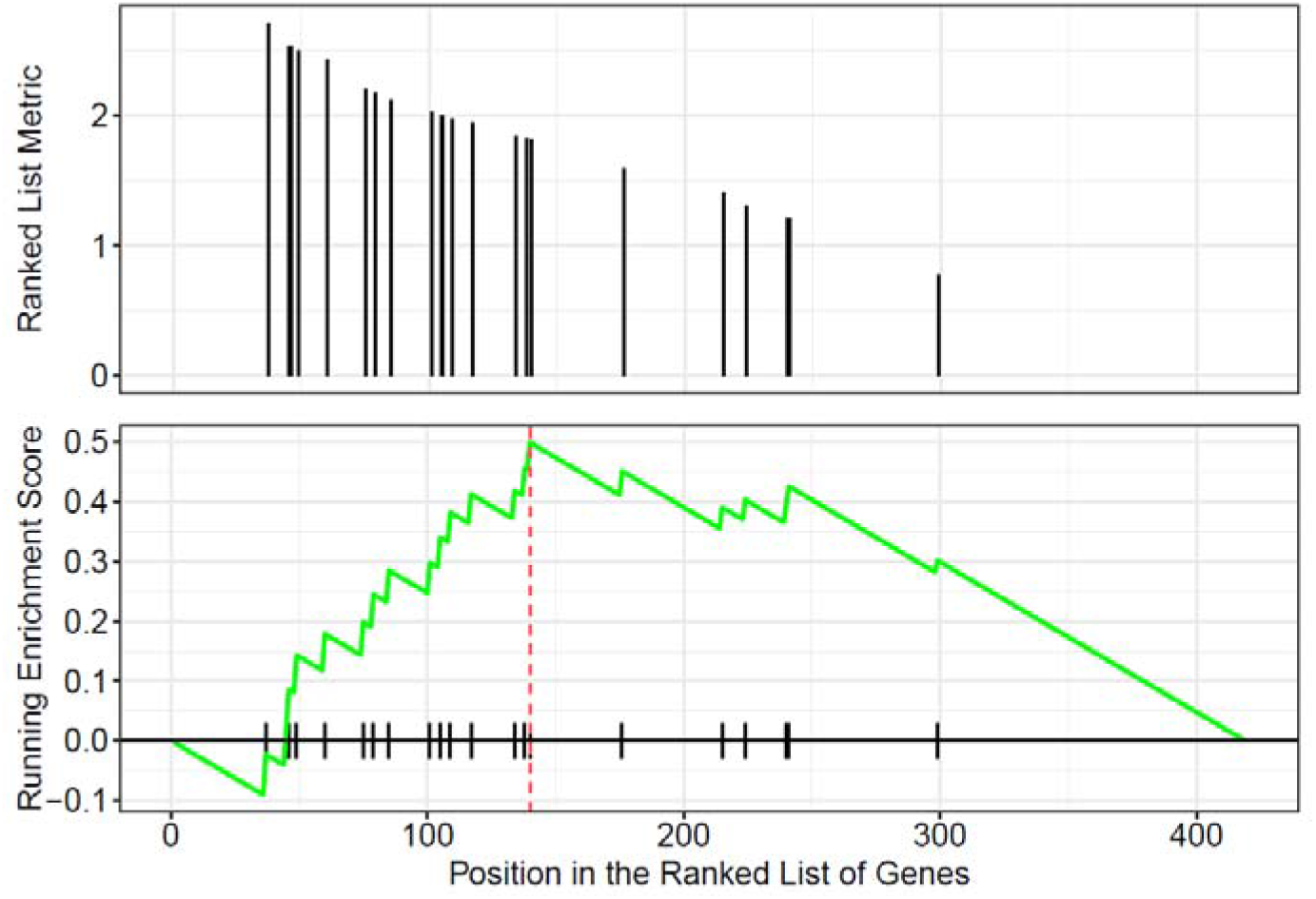
The KEGG plot of the cytokine-cytokine receptor interaction (hsa04060) pathway. The peak point of the green line shows over expression of the gene-set respect to the ranked list.

### 3.4 Gene-Drug Interactions

To explore the possible treatment options for younger MM patients, we performed a drug-gene interaction analysis of the top 20 significant DEGs with prescription drugs using DGIdb search tool. The name of drugs with associated genes are listed in Table 3.

**Table 3.**
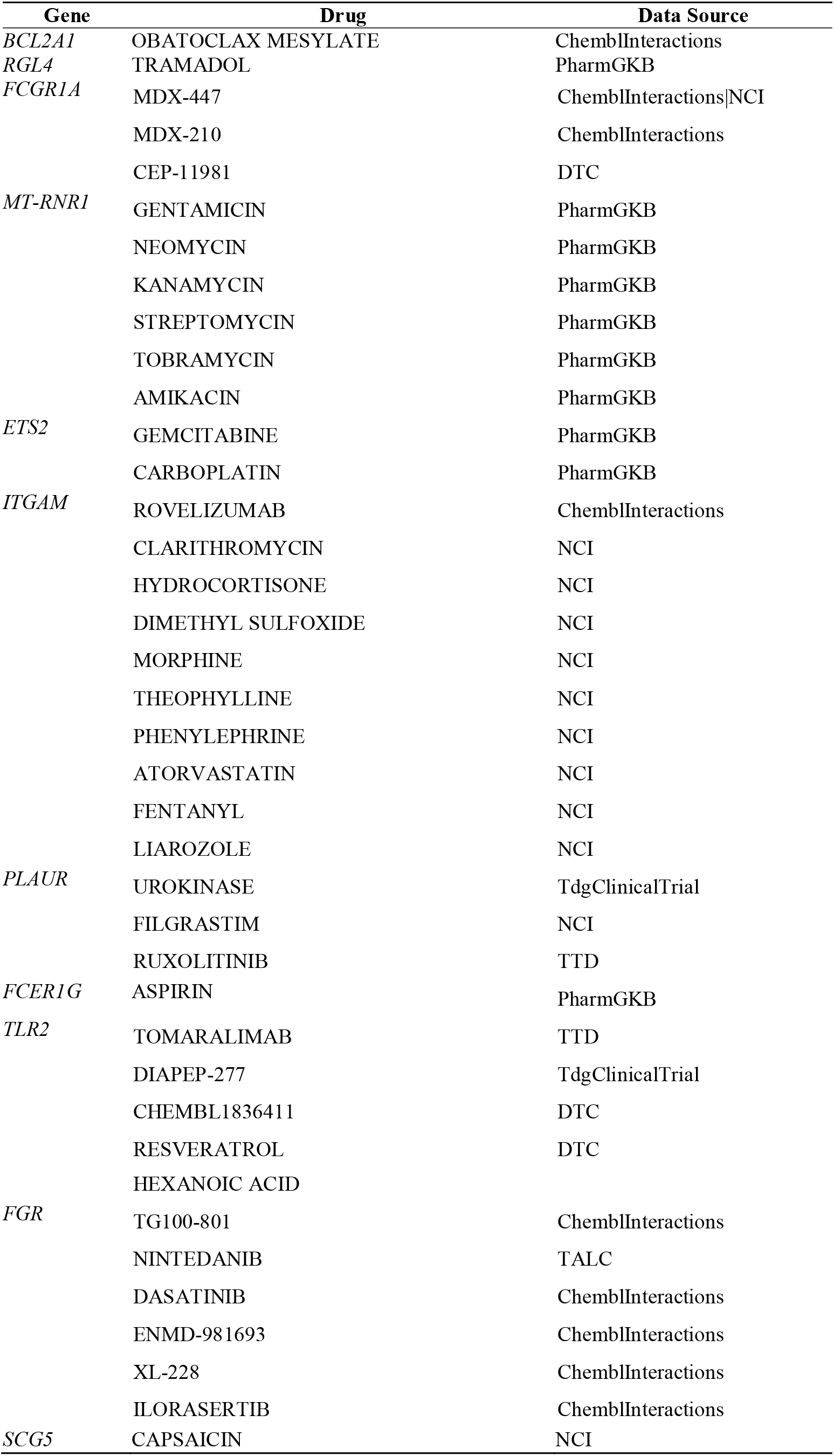
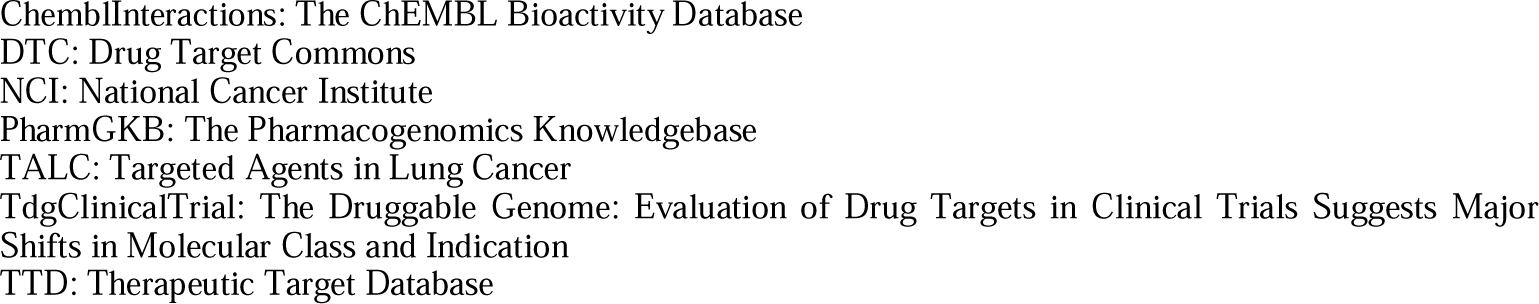
Gene-Drug Interaction Analysis Results.

## 4. Discussion

In this study, RNA-Seq data analysis of young and older patient groups showed that the DEGs in young patient group were mostly related to immune regulation. The only enriched pathway in GSEA was the “cytokine-cytokine receptor interaction”. Although these findings are not surprising considering age-related physiological immunity changes and the plasma cell origin of the disease, it is not yet known whether the immune pathway regulation changes are associated with different pathological processes in different age groups in MM. In a recent review review (Chatsirisupachai, Lagger, and de Magalhães 2022), age-associated molecular differences in various cancer types were examined indicating that immune-related pathways were enriched in different age groups in several cancers. A decline in T cell abundance by age, higher mutation burden, increased expression, and decreased promoter methylation of immune checkpoint genes were reported about immune regulation pathways. A better understanding of age-related immune regulation in MM could reveal which patients will benefit from specific immunotherapy.

Among the top 20 significant DEGs detected in this study, *FCGR1A, FCER1G and TLR2* were directly associated with immune-regulation.

*FCGR1A* encodes a protein that plays an important role in the immune response and it is highly expressed in various cancer types. In a recent study *FCGR1A* was found associated with infiltrating levels of CD4+ T cells, CD8+ T cells, B cells, macrophages, neutrophils, and dendritic cells in four cancers and suggested as a potential prognostic biomarker (Xu and Guo 2020).

*FCER1G* (Fc fragment Of IgE receptor Ig) is involved in the innate immunity. In a recent study, the expression level of *FCER1G* is found negatively correlated with myeloma progression, and high *FCER1G* expression suggested as a favorable biomarker in MM patients (Fu et al. 2020). Toll-like receptors (TLRs) are key immune receptors that recognize conserved molecular patterns expressed by pathogens and damaged cells. TLR activation causes inflammatory responses, modulation of cell cycle, apoptosis, or regulation of cell metabolism. In multiple myeloma, TLRs signaling is dysregulated and the expression of *TLR1, TLR2 and TLR6* seem to be higher in MM patients than in healthy donors and TLRs are suggested as future therapy targets in MM (Akesolo et al. 2022), (Thakur et al. 2015).

Furthermore, another DEG found in this study, *PGLYRP1*, encodes a protein that has peptidoglycan binding activity and involved in humoral immune response (Romanova et al. 2020).

Further investigation of the roles of these genes involved in immune regulation may shed light on innovations in the immunotherapy of the disease.

In addition to the genes involved in immune regulation, known proto-oncogenic genes (*BCL2, FGR*), genes under investigation for association with various cancers (*RGL4, MT-RNR1, ETS2, ENPP3, FUT7, NTNG2, PRAM1*) and an anti-angiogenic anti-tumoral gene (*PTX3*) were found in the top 20 significant DEGs in young patients.

B-cell lymphoma 2-related protein A1 (BCL2A1) is one of the anti-apoptotic BCL2 proteins, which is a target of nuclear factor jB (NF-jB) gene and it is expressed in the hematopoietic system in physiologic conditions. However, *BCL2A1* is overexpressed in a variety of cancer cells(Vogler 2012). In a recent study, it is reported that disease relapse in MM is related to increased NF_κ_B pathway activity, which causes increased *BCL2A1* expression (Spaan et al. 2021). Considering the upregulated *BCL2A1* expression in young MM patients, it can be hypothesized that *BCL2A1* may be a target for treatment with the current anti-cancer drug obatoklax mesylate currently used in acute myeloid leukemia and lymphoma.

The proto-oncogene *FGR* belongs to the Src family of tyrosine kinases; in physiological conditions it is expressed in hemopoietic cells. *FGR* overexpression promotes the development and rapid progression of acute myeloid leukemia (AML), ovarian and colorectal cancers (Perez et al. 2022).

In the drug-gene interaction analysis conducted in this study, drugs related to the pathways regulated by *BCL2 and FGR* genes and used in the treatment of various cancer types were identified. Further research is needed to understand the possible benefits of using these drugs in young MM patients.

*RGL4*, is known to be a regulator of the Ras pathway involved in cell proliferation. Although it has been suggested to contribute to the development of some T-cell malignancies (Leonardi et al. 2002), its role in certain cancer types and multiple myeloma has not been elucidated.

*MT-RNR1* over-expression can be associated with increased mtDNA levels in malignant transformation of myeloma cells. But its role in pathogenesis and as a possible treatment target remain unclear (Ruiz-Heredia et al. 2021).

*ETS2* is a downstream effector of the RAS/RAF/ERK pathway, which plays a critical role in cell proliferation also in the development of tumors. Its over-expression is reported as a poor prognostic factor in Acute Myeloid Leukemia (Fu et al. 2017).

*ENPP3* (ectonucleotide pyrophosphatase/phosphodiesterase 3) is known to be expressed in renal tubules, activated basophils and mast cells. Among cancers, it is expressed by renal cell carcinoma cells and anti-ENPP3 therapy is being investigated (Thompson et al. 2018).

*FUT7* is related to e-selectin ligand synthesis and its over-expression in MM was found to be related to significant inferior progression-free survival (Natoni et al. 2017).

*NTNG2* gene encodes a membrane binding netrin family protein, reported as related to various cancers and is shown as potential new tumor marker and therapeutic target (Hao et al. 2020) but its role in multiple myeloma pathogenesis has not been elucidated yet.

*PRAM1* encodes an adaptor protein is expressed and regulated during normal myelopoiesis. While it is known to be related to promyelocytic leukemia, it is not considered as a general oncogene (Moog-Lutz et al. 2001).

There is a need to investigate the effects of expression differences of these genes in MM, which have various roles in various cancers.

Only one gene, PTX3, was found in the top DEGs with anti-tumoral effects. PTX3 (Long Pentraxin 3) encodes a natural FGF antagonist that causes an anti-angiogenic effect. Moreover, an increased FGF/PTX3 ratio occurs during MM evolution. In a recent study, PTX3 overexpression was found associated with significantly reduced tumor burden in MM models (Ronca et al. 2021).

## 5. Conclusions

In this study, by comparing the transcriptome data of young and old MM patients, differentially expressed genes with oncogenic and immunological effects were identified. Replication of these research findings with independent data and supporting the information about the differentially expressed genes with functional studies may shed light on new treatment strategies.

## 6. Declarations

### Ethical Approval and Consent to participate

Not applicable for studies not involving humans

### Human and Animal Ethics

Not applicable for studies not involving humans and animals

### Consent for publication

All authors have read and agreed to the published version of the manuscript.

### Availability of supporting data

The MMRF CoMMpass (NCT01454297) datasets downloaded and analyzed for the current study are available through https://research.themmrf.org upon request. Source files used for this study are “MMRF_CoMMpass_IA17_star_geneUnstranded_counts.tsv” and “MMRF_CoMMpass_IA17_PER_PATIENT_V2.tsv” access date: 1 July 2022.

### Authors’ contributions

M.K. Conceptualization, methodology, formal analysis, investigation, resources, data curation, writing original draft preparation, visualization,

## Acknowledgements

All data in this study were generated as part of the Multiple Myeloma Research Foundation Personalized Medicine Initiatives (https://research.themmrf.org and www.themmrf.org). The author thanks Sema Sirma Ekmekci (Istanbul University Aziz Sancar Institude of Experimental Medicine Department of Genetics) and Beyza Oluk (Istanbul University Medical Faculty Department of Hematology for encouraging work on multiple myeloma.

## Supplementary Materials

Supplementary Data 1: List of the All Significant DEGs; Supplementary Data 2: List of the All Significant GO Terms.

